# Computational methods for the discovery and annotation of viral integrations

**DOI:** 10.1101/2021.08.28.458009

**Authors:** Umberto Palatini, Elisa Pischedda, Mariangela Bonizzoni

## Abstract

The transfer of genetic material between viruses and eukaryotic cells is pervasive. Somatic integrations of DNA viruses and retroviruses have been linked to persistent viral infection and genotoxic effects. Integrations into germline cells, referred to as Endogenous Viral Elements (EVEs), can be co-opted for host functions. Besides DNA viruses and retroviruses, EVEs can also derive from nonretroviral RNA viruses, which have often been observed in piRNA clusters. Here, we describe a bioinformatic framework to annotate EVEs in a genome assembly, study their widespread occurrence and polymorphism and identify sample-specific viral integrations using whole-genome sequencing data.

## 1. Introduction

Lateral gene transfer (LT) from viruses to eukaryotic cells is a well-recognized phenomenon [1]. Somatic integrations of DNA viruses and retroviruses have been linked to persistent viral infection and genotoxic effects, including various types of cancer [2, 3]. Viral sequences that integrate into germline cells can be transmitted vertically and co-opted for host functions. These Endogenous Viral Elements (EVEs) have long been known, with studies focusing mainly on EVEs from retroviruses in mammalian genomes [4].

In the last decade, innovative Next Generation Sequencing (NGS) technologies paved the way to produce high quality genome assemblies for non-model eukaryotes [5]. This genomic leap led to the discovery of EVEs not only from DNA viruses and retroviruses, but also from nonretroviral RNA viruses in organisms of different evolutionary lineages, including arthropods (i.e. ticks, mosquitoes, bees, crustaceans), fish, snakes, birds, vertebrates (i.e. primates, mouse, rat, opossum) and plants [6–9]. These nonretroviral (nr) EVEs are highly variable in number and many occur proximal to transposable element (TE) sequences in piRNA clusters [7, 10]. Experimental evidence is increasing on the role of nrEVEs in different biological processes such as antiviral immunity, tolerance to cognate viral infection and regulation of host gene expression [6]. However, the enrichment of nrEVEs in repetitive piRNA clusters complicates their genome annotation and hinders the ability to detect viral integrations through alignment to a reference genome of whole genome sequencing (WGS) data from natural samples or samples collected under hypothesis-driven conditions. The commands and pipelines we describe here were designed to overcome these issues and allow the precise annotation of nrEVEs in a genome assembly and to study of their sequence polymorphisms and distribution. Because the situation of nrEVEs is the most complex, our computation framework can be adopted to discover and annotate any viral integration, providing an *ad hoc* viral database to interrogate.

Our bioinformatic protocol accomplishes three different tasks:

1. Find and annotate EVEs in a genome assembly.
2. Test EVE frequency and their sequence polymorphism using WGS data.
3. Identify, from WGS data, viral integrations that are absent in the reference genome assembly. The execution of task 1 depends on BLASTx, a popular algorithm for the comparison of proteins to nucleic acids [11]. BLASTx is used to identify sequences in a genome assembly showing similarity to a user-provided set of viral proteins. The resulting BLASTx hits are further filtered by custom scripts (see 3.1) to reduce instances of false positive hits. Tasks 2 and 3 are implemented with WGS short reads that are mapped to a genome assembly. Task 3 is executed by running Vy-PER [12], a pipeline originally designed to identify viral integrations into the human genome, followed by ViR [13], a pipeline designed to improve predictions of new integration sites by accounting for dispersion of reads in repetitive DNA sequences, such as piRNA clusters. The identification of viral integrations using alignment of WGS data to a genome assembly is based on the identification of chimeric reads (a read pair in which one read maps to the host genome, hereafter referred to as host read, and the other read to a virus, hereafter referred to as viral read) and/or unmapped reads, which are extracted from the WGS dataset. The three tasks described above can be executed with any genome assembly as long as a fasta file is available, without the need for gene models.

## 2. Materials

The framework we describe here is divided into three procedures that can be used to answer different questions about the presence of EVEs in an organism’s genome. Each procedure can be run independently depending on the user’s needs. The annotation of existing EVEs in a reference assembly (obtained with procedure 1) is strictly required for procedure 2 and is advised before running procedure 3. Performing each of the three parts of this protocol requires different starting data, programs, and computational power. Familiarity with the Unix environment and command-line and the ability to understand and run simple scripts is advised. Users should also be familiar with NGS and have a clear idea of the theory behind short reads sequencing technologies, as well as experience with BLAST and BWA [11, 14].

### 2.1 Required programs and scripts

Our protocol can be run entirely with open-source tools and scripts. A Unix machine or cluster is required to install and run all the programs (see 2.2). The programs required for each step of the protocol are listed below. For the installation process and additional dependencies refer to the guides of each tool. Some of these tools/pipelines necessitate both Python 2 and 3 (https://www.python.org/download/releases/) to be run. It is convenient to install a program for the manipulation of fasta sequences with a graphical user interface and the Integrative Genome Browser (IGV) [15], or a similar software, to visualize alignments of sequencing reads.

#### 2.1.1. Procedure 1, annotation of EVEs in a genome assembly

1. BLAST + [16] (https://ftp.ncbi.nlm.nih.gov/blast/executables/blast+/LATEST/) or DIAMOND [17] (https://github.com/bbuchfink/diamond
2. EVE_finder [18] (https://data.mendeley.com/datasets/d6zf6fvzwn/1)
3. Refine EVEs annotation pipeline (https://github.com/BonizzoniLab/Refine_EVEs_annotation)
4. BEDtools [19] (https://bedtools.readthedocs.io/en/latest/)
5. Virus-Host Classifier [20] (https://github.com/Kzra/VHost-Classifier)
6. Taxonkit [21] (https://bioinf.shenwei.me/taxonkit/)

#### 2.1.2. Procedure 2, Assessment of EVEs polymorphism

1. GATK [22] (https://gatk.broadinstitute.org/hc/en-us)
2. Platypus [23] (https://www.well.ox.ac.uk/research/research-groups/lunter-group/lunter-group/platypus-a-haplotype-based-variant-caller-for-next-generation-sequence-data)
3. Freebayes [24] (https://github.com/freebayes/freebayes)
4. VarDict [25] (https://github.com/AstraZeneca-NGS/VarDict)
5. BCFtools [26] (http://www.htslib.org/download/)
6. SVD pipeline [27] (https://github.com/BonizzoniLab/SVD)

#### 2.1.3. Procedure 3, Identification of novel EVEs

1. Vy-PER [12] (https://www.ikmb.uni-kiel.de/vy-per/).
2. ViR [13] (https://github.com/epischedda/ViR)
3. BLAST+ [16] (https://ftp.ncbi.nlm.nih.gov/blast/executables/blast+/LATEST/)
4. BEDtools [19] (https://bedtools.readthedocs.io/en/latest/)
5. BWA [14] (http://bio-bwa.sourceforge.net/bwa.shtml)
6. BLAT [28] (http://hgdownload.soe.ucsc.edu/admin/exe/)
7. Phobos 3.3.12 (Christoph Mayer, http://www.rub.de/spezzoo/cm/cm_phobos.htm)
8. SAMtools [29] (http://www.htslib.org/download/)
9. Trinity [30] (https://github.com/trinityrnaseq/trinityrnaseq/)

### 2.2. Computational power

The most important factors influencing the computational power, running time and disk occupancy required by the procedures described here are 1) the size of the reference genome assembly, 2) the size and number of the raw sequencing reads (fastq) files and the consequent aligned reads (sam/bam) files, and 3) the size of the databases (of viral proteins and/or genomes) used to search for EVEs. A high-performance computing (HPC) cluster or a dedicated server with at least 32 computing threads and 64GB of RAM are recommended, especially to run Vy-PER as this pipeline can run multiple alignment processes in parallel via a scheduling manager [12, 13]. The advantage of running multiple alignments in parallel depends on the behavior of BWA, the most computationally-intensive step in the protocol. BWA run time begins to stabilize at about 10-12 threads when aligning a single sample [31]. This means that increasing further the number of threads does not significantly increase the “wall clock” time required to execute the program. Splitting the WGS data (fastq files) into multiple parts and running them separately is the best solution to maximize the outcome of having multiples of 8 threads available.

### 2.3 Notes before beginning

1. Carefully read the manuals of the tools described in this protocol. Detailed information on the execution of the programs and their options are available online. Additionally, Vy-PER and ViR include step-by-step descriptions and example files that can be used to test the configuration before using actual data [12, 13].
2. Commands to execute the three tasks described here are not concatenated to make it easier for users to understand each passage. However, they can be concatenated together by preparing a script or integrating them into a workflow management tool. Commands/scripts can be run in the background using the nohup command or a HPC cluster workload manager.
3. The design of the database of viral sequences for the annotation of EVEs in a reference genome (procedure 1) and for the search of novel EVEs (procedure 3) should be carefully considered, as it varies depending on the research question, the desired sensitivity, and the available computational power.

## 3. Methods

### 3.1 Annotation of EVEs in a genome assembly

Among the different strategies that have been proposed for EVEs annotation [7, 10, 18, 32–35], we consider the one described below as the best compromise between sensitivity, precision and computing time. Any reference genome assembly can be screened for viral integrations using an approach based on the BLASTx algorithm included in the BLAST+ suite [16]. BLASTx automatically translates query nucleotide sequences and compares them against a target database of proteins using a heuristic approach which initially finds short matches between two sequences (seeding). After BLASTx-based identification of potential EVEs [11, 18], automatic filtering steps are implemented to reduce false positive results. Filtering relies not only on a “reverse” BLASTx search of the BLASTx hits against the entire NCBI protein nr database but also on a taxonomic analysis of each putative EVEs to eliminate results of non-viral origin.

#### 3.1.1 Preliminary operations

The following items are required before launching any program:

- A DNA fasta file of the genome assembly to be screened. The names of the DNA sequences in the fasta file should not contain spaces or special characters and should be concise to reduce the risk of errors in downstream applications.
- A database of viral proteins in fasta format. Be careful not to include duplicate sequences.
- A large reference protein database to eliminate false positive results. These databases are large (up to 200GB) so ensure to have enough space on the working folder.

The size of the target DNA file and the protein databases will influence both the time required for the BLASTx run and the size of the output file. A regular update of the taxonomic databases used by Taxonkit and VHost-Classifier is required to classify predicted EVEs according to the most recent taxonomy [20, 21]. The update can be done following instructions included in the readme file of the EVEs annotation refinement pipeline (https://github.com/BonizzoniLab/Refine_EVEs_annotation). Of note, Taxonkit requires the taxon database to be in the folder “home/user/.taxonkit” by default. Refer to the tool manual (https://bioinf.shenwei.me/taxonkit/) for more information.

#### 3.1.2 Running the pipeline

1. If an indexed database is not already available, create one using the command included in the BLAST+ package:

~~~
$ makeblastdb -in ProteinDatabase.fasta -dbtype prot
~~~
2. After building the database files, run BLASTx using as query the target assembly:

~~~
$ blastx -query ReferenceGenome.fasta -db ProteinDatabase.fasta -evalue 1e-6
-num_threads 8 -outfmt ‘6 qseqid qstart qend salltitles evalue qframe pident qcovs sstart
send slen’ -out EVEs_target.blastx
~~~ The “-evalue” option defines a threshold for the hits on the basis of the Expect Value. The closer to zero the E-Value threshold is set, the more significant the retained hits will be. The “-num_threads” option sets the number of computing threads and should be adjusted based on your system and data. For genome assemblies with a size larger than 1Gbp, we suggest using 8 threads, as increasing further the number of threads does not increase the computing speed. Threads can be reduced for smaller assemblies or protein databases or if extended computing time is not an issue.
3. Run the “Blast_to_Bed3.py” script in the eve_finder toolbox to convert the tabular BLASTx output to Browser Extensible Data (BED) format. The output will have the “EVEs.bed” suffix:

~~~
$ python eve_finder/Blast_to_Bed3.py EVEs_target.blastx
~~~ If the script produces an error, try adding an underscore (“_”) in the BLASTx output file name.
4. Sort the resulting BED file for position relative to the target assembly and merge overlapping hits, if present:

~~~
$ bedtools sort -i EVEs_target.EVEs.bed > EVEs.sorted.bed
$ bedtools merge -i EVEs.sorted.bed -c 4,5,6,7,8,9,10,11 -o
collapse,collapse,distinct,collapse,collapse,collapse,collapse,collapse > EVEs.merged.bed
~~~
5. Use the “Top_score_BED2.py” script in the “eve_finder” toolbox to select the best hit from each cluster of overlapping hits. The output is a bed file containing the position of the uniquely selected EVEs in the reference assembly.

~~~
$ python EVE_finder/Top_score_BED2.py EVEs.merged.bed EVEs_top.bed
~~~
6. Get the fasta sequence of the EVEs from the genome assembly. Each EVE will be named with a combination of the best hit viral species and starting genomic position:

~~~
$ bedtools getfasta -s -name -fi ReferenceGenome.fasta -bed EVEs_top.bed -fo EVEs_top.fasta
~~~
7. Use the final collection of EVE sequences extracted from the assembly as a query for a BLASTx against a large Database of proteins. We suggest using DIAMOND [17] for this search because it is faster than the native BLASTx algorithm in the BLAST+ [16] package when used to align queries on a large database. This reverse BLASTx can be done against any database of choice but, to our knowledge, the most comprehensive and up to date is the NCBI protein nr database (https://www.ncbi.nlm.nih.gov/refseq/about/nonredundantproteins/), which includes protein sequences from NCBI RefSeq, GenPept, Swissprot, PDB and other databases. When using DIAMOND [17], remember to build your database from the nr database fasta file (https://ftp.ncbi.nlm.nih.gov/blast/db/FASTA/nr.gz) with the “diamond makedb” command first and then launch the alignment with the “diamond blastx” command:

~~~
$ diamond makedb --in nr_database.fasta -d nr_diamond.dmnd
$ diamond blastx -d nr_diamond.dmnd -e 1e-06 --threads 8 -f 6 qseqid qstart
qend salltitles evalue qframe pident qcovhsp sstart send slen staxids -q
TopHits.fasta -o EVEsTop_nr.blastx
~~~ When using BLAST+ [16], download and extract the preformatted nr database (https://ftp.ncbi.nlm.nih.gov/blast/db/) and directly use the “blastx” command:

~~~
$ blastx -query EVEs.top.fasta -db nr/nr_protein -evalue 1e-6 -num_threads 8
-outfmt ‘6 qseqid qstart qend salltitles evalue qframe pident qcovs sstart
send slen staxids’ -out EVEsTop_nr.blastx
~~~
8. Run the custom script to select unique NCBI taxonomical IDs and extract hits with similarity to viruses from the BLASTx table. This step is important to remove false positives that do not have similarity to any known viral sequences, have a strong similarity to eukaryotic proteins and/or are low-complexity sequences (e.g., tandem repeats). In addition, this script assigns EVEs to a viral species.

~~~
$ bash Refine_EVE_Annotation.sh \
-pipeline_directory Refine_EVE_annotation_folder/ \
-tool diamond \
-VHC_directory VHost-Classifier/ \
-file_blastx EVEsTop_nr.blastx \
-file_bed_tophit EVEsTop.bed \
-output_directory Output_directory \
-taxonkit_exe Taxonkit_0.6/taxonkit
~~~ Change the “-tool” option to “diamond” or “blastx” depending on the chosen tool.

#### 3.1.2 pipeline output

Two folders are created in the path defined by the user. The “VHC/” folder contains the files created by VirusHost-Classifier while the “Output/” folder contains the tab separated value (tsv) tables produced by the pipeline. The “CompleteTable_classified” file in the “Output/” folder incorporates comprehensive data about the genomic position of each EVE, the best viral and non-viral BLASTx/DIAMOND hits, and the assignment of each EVE to the viral taxonomy. For a detailed description of the fields in the table, refer to the pipeline readme (https://github.com/BonizzoniLab/Refine_EVEs_annotation).

### 3.2 Assessment of EVE polymorphism

We developed the Structural Variants Definition (SVD) pipeline (Figure 1) to test the occurrence and sequence polymorphism of EVEs in samples collected from the field or under hypothesis-driven experimental conditions (https://github.com/BonizzoniLab/SVD). The SVD pipeline can be applied to WGS data from one or multiple samples, the latter called a population. The SVD pipeline relies on four Variant Callers (Freebayes [24], VarDict [25], GATK [22] and Platypus [23]) for the detection of Single Nucleotide Polymorphisms (SNPs) and Insertions and/or Deletions (INDELs) in the samples and it implements four subsequent steps (Figure 1): 1) coverage evaluation; 2) variant calling; 3) post variant calling processing and 4) final alleles determination.

**Figure 1.**
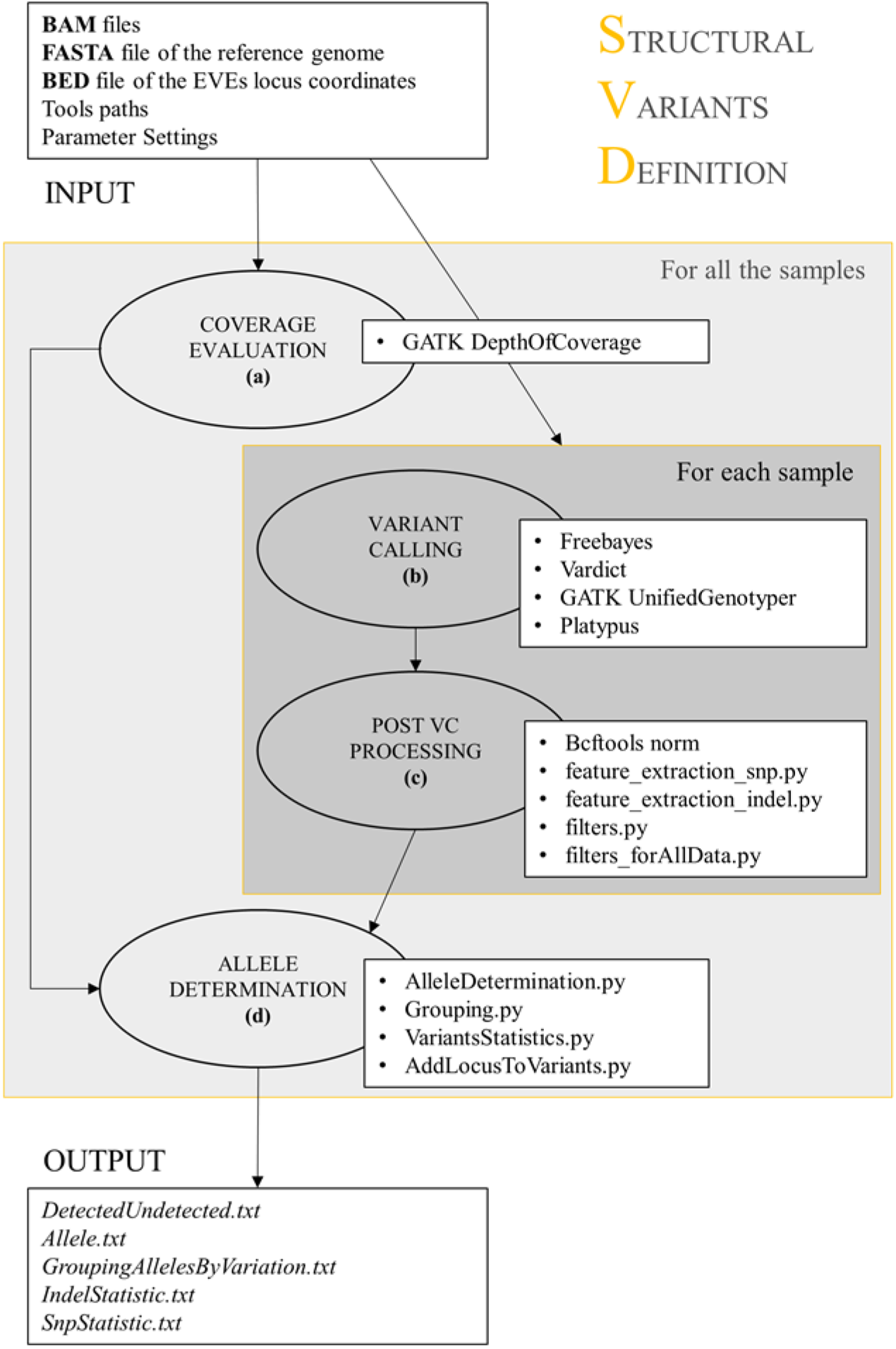

#### 3.2.1 Preliminary operations

The input files required by the SVD pipeline are:

- A reference genome assembly in fasta format.
- A bed file with coordinates of EVEs which have been annotated in the genome assembly. In addition, Platypus [23] requires a file where each feature is annotated with the format: “scaffold:start-end”. Refer to the documentation of the program for additional information.
- For each sample, a bam file containing the WGS reads aligned to the reference genome assembly. A list containing the full paths of all the bam files must be prepared.

#### 3.2.2 Running the SVD pipeline

To perform this procedure, WGS data must be aligned on a reference genome using BWA [14] or a similar short reads aligner to produce bam files. Here we will show the alignment of paired end reads from an individual using bwa mem.

1. Align the paired end reads with BWA and sort them with SAMTools. Here we concatenate the commands and use default parameters.

~~~
$ bwa mem -t 16 -R RGID reference_genome_assembly.fasta R1.fastq R1.fastq | samtools view -@ 8 -m 8G -b Sample.bam
~~~
2. Sort the bam file.

~~~
$ samtools sort -@ 8 -m 8G Sample.bam -o Sample.sorted.bam
~~~
3. Run the SVD pipeline through a single command line where all the input files and programs can be set. The user can define generalized parameters to filter variant calling results. The pipeline will use these parameters to homogenize the identified variants in terms of features common to the different variant callers [27]. The following command runs the pipeline with default parameters. Refer to the program manual for detailed information about the options (https://github.com/BonizzoniLab/SVD).

~~~
$ bash StructuralVariantDefinition.sh \
-c samples_BAM_files.list \
-b reference_EVEs.bed \
-b_pl platypus_EVEs.txt \
-f reference_genome_assembly.fasta \
-i StructuralVariantsDefinition_09_11_18/ \
-o output_folder/ \
-fbpath freebayes/bin/freebayes \
-gkpath gatk.jar \
-vdpath VarDict/vardict
--fileR VarDict/teststrandbias.R
--filePl VarDict/var2vcf_valid.pl \
-plpath Platypus/bin/Platypus.py \
-btpath bcftools/bcftools \
-th 4 --ram 12g --MIN_MQ 20 --MIN_BQ 20 --MIN_AF 0.1 --MIN_AO 2 --MIN_COV 8 -
-MAX_DEPTH 5000 --DP_expected_mean 20 -dp 8 -af 0.1 --AFallData 0.1 --
MaxStrBias 0.01 --MinLengthINDELallData 6 --NumCallersallData 2 --
minReadsAllDef 5 --minLengthAllele 30 --thresholdSimilarity 0.05
~~~

#### 3.2.3 SVD pipeline output

The pipeline produces the following outputs:

- A list of the annotated EVEs that are also present in the analyzed samples. The frequency of EVE occurrence in the sample(s) can be calculated from this information.
- For each sample, a summary table showing the number of variants found in each locus with the predicted zygosity (homo- or heterozygous) in the sample plus additional information on each allele (e.g., number of reads supporting the variant or reference allele, their orientation and mapping quality, which callers identified the variant). For each EVE, a list of detected SNPs and INDELs and a list of different alleles found across all analyzed samples.

Refer to the manual of the pipeline (https://github.com/BonizzoniLab/SVD) for more information about the output fields.

### 3.3 Identification of novel EVEs

When WGS data are available, it is possible not only to study the polymorphism of EVEs that are annotated in the reference genome assembly, but also to search for sample-specific EVEs that are not present in the reference assembly. Hereafter we will call the latter as novel EVEs. Several pipelines have been developed in the context of cancer genetics to identify novel viral integrations (for a review see [36]). Among these pipelines, Vy-PER (Virus integration detection by Paired End Reads) [12] emerges for its speed, accuracy of predictions and for the possibility to test integrations derived from more than one viral species simultaneously. The discovery of viral integrations from different viral species in several non-model organisms whose available genome assemblies are often still fragmented, and the enrichment of nrEVEs in repetitive DNA, including piRNA clusters, increases the complexity of EVE identification because reads supporting an EVE could be scattered across repetitive regions. To solve this issue, we developed ViR (Figure 2) [13].

**Figure 2.**
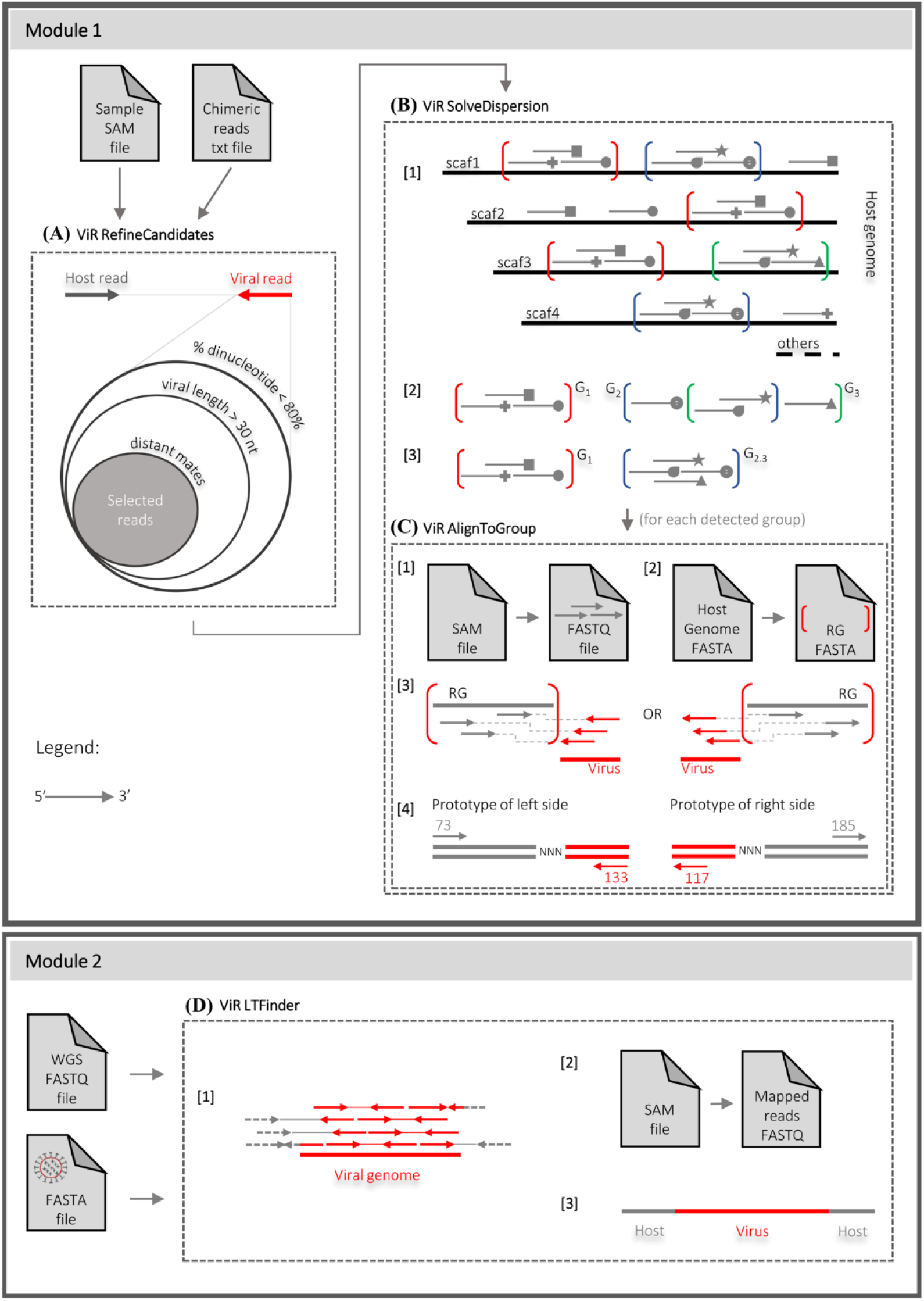

Vy-PER takes advantage of paired-end reads that are first aligned to the reference assembly using BWA [12]. Chimeric read pairs, in which one read maps to the reference genome and the other one does not, are selected. In these pairs, the read mapping to the reference genome is referred to as host read. Among the selected unmapped reads, low-complexity reads are discarded using Phobos [37]. The remaining unmapped reads are aligned with BLAT [28] to a user-defined viral genome database. Only the top 3 virus candidates per integration site are retained. Finally, a script refines the output including the final virus integration candidates. ViR works downstream of Vy-PER, or any other paired end reads-based EVE prediction algorithm, to improve predictions of viral integration sites by addressing the dispersion of reads due to intrasample variability (Figure 2) [13]. Intrasample variability can be due to repetitive DNA and/or fragmentation in the assembly. ViR is composed of four scripts, which work in two modules (Figure 2). The first module contains three scripts that help overcoming the dispersion of host reads by grouping together reads that map to regions of the genome with the same sequence, defined as “read groups”. The second module includes a single script, “ViR_LTFinder.sh”, designed to test for an integration from non-host sequences which have a user-defined percentage of similarity to host sequences. This script can assemble a consensus sequence extending the viral integrations if there are enough unique reads supporting the assembly. Vy-PER output files include a table of the top 10 virus candidates, a table of the clusters (genomic windows, number of candidates, virus name and NCBI ID), a detailed table of unfiltered virus candidates and a fasta file for each virus candidate for optional manual alignment/checking. ViR can be integrated with additional information such as 1) a list of the annotated EVEs, to correlate novel EVEs with the ones in the reference assembly; 2) a list of piRNA clusters and TEs to assess the genomic context of novel EVEs.

#### 3.3.1 Preliminary operations

The input files required by Vy-PER are:

- A reference genome assembly in fasta format.
- Two fastq files for each sample containing the raw paired end reads. Normally these files have the R1 and R2 suffixes, respectively. Reads should be clean from adapters and low-quality bases should have been trimmed.
- A database of viral genomes or sequences in fasta nucleotide format that will be used by BLAT to select reads of viral origins. Notice that this must be a nucleotide database unlike the one previously used to annotate EVEs, which was a protein database.

ViR requires:

- A reference genome assembly.
- The output from Vy-PER, more specifically:

□ A sam file containing reads aligned on the target genome assembly
□ A tab separated table of the chimeric reads identified by Vy-PER. The file containing the chimeric reads can be prepared directly from the output of Vy-PER and an example can be found in the”/ViR examples” folder in the ViR download.

In case sam files from multiple samples must be analyzed with ViR, a list of the sam files full paths must be provided in a text file.

Optional files can be provided to ViR. These files are:

- A bed file with the coordinates of the annotated EVEs in the reference assembly
- A bed file of the piRNA clusters annotated in the reference assembly
- A fasta file containing the nucleotide sequences of predicted TEs

#### 3.3.2 Running Vy-PER

As stated above, Vy-PER is computationally demanding, and it is a good idea to run it on a cluster [12]. With this idea in mind, Forster et al. split Vy-PER into several scripts to allow the user to optimize the number of cores used to run each script and implement a per-lane processing approach that results in better parallelization and faster run time. The Vy-PER download (www.ikmb.uni-kiel.de/vy-per/) includes example scripts for a Linux cluster that can be easily reconfigured for any Unix machine. For the final filtering step these scripts use a Smith-Waterman implementation on the FPGA-based system RIVYERA (www.sciengines.com), but for general users, Vy-PER can use BLAT on a Linux cluster. We suggest building Vy-PER with a tool or library for parallelized workflow management for high-throughput analysis such as Cosmos [38], uap [39] or HaTSPiL [40]. For clarity, we will describe here the individual Vy-PER scripts and their usage.

1. Align the WGS reads to the host genome using “BWA aln” on each fastq file and “BWA sampe” to generate the paired end sam file:

~~~
$ bwa aln ReferenceGenome.fasta -n 2 -q 15 -l 5000 -t 8 R1.fastq > R1.sai
$ bwa aln ReferenceGenome.fasta -n 2 -q 15 -l 5000 -t 8 R2.fastq > R2.sai
$ bwa sampe ReferenceGenome.fasta R1.sai R2.sai R1.fastq R2.fastq -a 500 > AllReads.sam
~~~ The “-a” option in the “bwa sampe” command line defines the maximum insert size and should be changed according to the insert size of the sequenced DNA library. Read groups (RG) can be added using the -r option.
2. Convert the output SAM file into a sorted bam file. If a fai index for the reference genome is not present, create it using the “samtools faidx” command.

~~~
$ samtools view -@ 4 -u -h -b -T ReferenceGenome.fasta.fai AlignedSample.bam
| samtools sort -@ 4 > SortedReads.bam
~~~
3. Extract the unmapped read whose mate is mapped on the reference genome to a new sam file, indicating that the original fragment partly matched the reference genome.

~~~
$ samtools view -f 4 -F 264 > UnmappedReads.sam
~~~
4. Convert the unmapped reads sam file to fasta using the first script of the Vy-PER package.

~~~
$ Vy-PER_sam2fas_se UnmappedReads.sam UnmappedReads.fasta
~~~
5. Identify reads with less than 30 bp (or a user-selected value) of non-repetitive DNA with Phobos [37] and use the “Vy-PER_sam2fas_se.py” script again to only extract these reads.

~~~
$ phobos --outputFormat 3 UnmappedReads.fasta phobosReads.fp3
$ Vy-PER_sam2fas_se -fp3 phobosReads.fp3 30 UnmappedReads.sam UnmappedReads_passed.fasta
~~~
6. Run BLAT to compare the extracted reads passing the Phobos filtering to the fasta database of viral genomes. BLAT also requires a 2bit database index that can be created with the “faToTwoBit” executable in the blat suite [28]. Create the 2bit files for the viral genomes database and for the reference assembly fasta file (it will be used by the last Vy-PER script).

~~~
$ blat -out=blast8 -noTrimA -t=dna -q=dna -maxGap=0 -fastMap
ViralDatabase.2bit UnmappedReads_passed.fasta UnmappedReads_blat.txt
~~~
7. Run the “Vy-PER_blatsam.py” script to extract the reads mapping to viruses from the original paired-end SAM files (these are the novel EVEs) and their mates mapping to the reference genome assembly. A summary table that shows EVEs sorted by their predicted genomic position and the three best virus candidates for each integration site is produced as output.

~~~
Vy-PER_blatsam SampleID ViralDatabase.fasta UnmappedReads_blat.txt
UnmappedReads_passed.fasta ReferenceGenome.fasta AllReads.sam
VyPER_OutputTable.txt FastaFolder/
~~~
8. Run the script “Vy-PER_final_filtering.py” to refine the results and to get the final output. This script removes EVE candidates whose sequence is for more than 50% repetitive DNA and filters the remaining reads to define host/virus chimera clusters.

~~~
Vy-PER_final_filtering -p 1000 10 10 0.01 0.5 0.95 3 0.90 0 swout.txt 0
VyPER_OutputTable.txt ReferenceGenome.2bit ReferenceGenome.fasta.fai SampleID
~~~ The “-p” option defines the parameters for clustering the host/virus chimeras and can be adjusted to obtain more or less stringent results. The “swout.txt” file will be empty if no FPGA-based Smith-Waterman [41] is implemented as in this example.

#### 3.3.3 Vy-PER output

The pipeline produces the following output files:

- A table of the top 10 virus candidates.
- A table of the clusters (including these fields: genomic windows, number of candidates, virus name and NCBI ID).
- A detailed table of unfiltered virus candidates.
- FASTA files for each virus candidate for optional manual alignment/checking.

In addition, the “rscript_ideogram.R” script can be used to plot in R a summary graph in PDF for the EVEs found by Vy-PER if the hg19 human reference genome assembly (www.ncbi.nlm.nih.gov/assembly/GCF_000001405.13/) was used.

#### 3.3.3 Running ViR module 1

The first module of ViR [13] is run on results from Vy-PER [12] or any similar program for the identification of host/virus chimera from paired-end reads. The module is made by three separate scripts which refine Vy-PER predictions and overcome the dispersion of reads due to intrasample variability.

1. Run the “ViR_RefineCandidates.sh” script to select the best candidate pairs from a list of host/virus chimeras supporting a novel EVE. By default, the script excludes viral reads that do not satisfy all of these conditions: 1) the viral portion in the read is shorter than 30bp; 2) complex and repetitive nucleotides represent more than 80% of each read content and 3) Read mates do not align together within a 10 kb window in the reference genome. These settings can be modified by the user.

~~~
$ bash VIR-master/ViR_RefineCandidates.sh \
-work_files_dir VIR-master/ \
-sample_name SampleID \
-sam_file AllReads.sam \
-chimeric_reads_file sample_chimeric_reads.txt \
-out output_directory_refineCandidates/ \
-reference_fasta ReferenceGenome.fasta \
-path_to_blastn blastn \
-path_to_bedtools bedtools \
-max_percentage_dinucleotide_in_ViralSeq 0.8 -minimum_virus_len 30 - blastn_evalue 1e-15 -min_mate_distance 10000
~~~ The outputs are a table containing the chimeric reads passing the filters and the fasta sequences of the viral and host reads.
2. Run the “ViR_SolveDispersion.sh” script to solve the dispersion of host reads by grouping together reads that map to regions of the genome with the same sequence (Read Groups). Reads mapping to regions of the genome with the same sequence (hereafter called equivalent regions) are identified and merged to define Read Groups. The input for this script is the output folder of the previous script, the reference genome, and a list of samples to be analyzed together. In addition, a fasta file of known transposable element sequences in the species and bed files for known EVEs and piRNA clusters can be provided to correlate the novel EVEs with these genome features. Running this script will automatically launch the third script, “ViR_AlignToGroup.sh”.

~~~
$ bash VIR-master/ViR_SolveDispersion.sh \
-work_files_dir VIR-master/ \
-outdict_RefCand output_directory_refineCandidates/ \
-analysis_name SAMPLE_ID \
-sample_list sample_list.list \
-out output_directory_solveDispersion/ \
-reference_fasta ReferenceGenome.fasta \
-repreg_fasta Transposable_elements.fa -min_TE_al_length 100 \
-bed_EVE_annotated EVEs.bed -eve_dist 10000 \
-bed_piwi_clusters piwiClusters.bed -piwi_dist 0 \
-path_to_blastn blastn \
-path_to_bedtools bedtools \
-trinity_exe Trinity \
-samtools_exe samtools \
-bwa_exe bwa \
-merge_dist 1000 -minReads_inRegion 2 -percReadsShared_inGroup_union 0.8
~~~

#### 3.3.4 ViR module 1 output

Five files and a directory “RG/” are produced. The files include detailed information and fasta sequences of the read groups identified in the samples and of all the possible equivalent regions in which they may be positioned in the genome. Also, information about the integration sites and the related boundaries regions are provided. The “RG/” directory includes the realignment of the reads in their relative assigned read group and consensus sequence for the integrations, if enough data was available. A detailed description of the output files can be found in the ViR readme (https://github.com/epischedda/ViR).

#### 3.3.4 Running ViR_LTFinder.sh

The second ViR module contains a single script: “ViR_LTFinder.sh”. The script maps WGS reads to a selected non-host fasta sequence using BWA and mapped reads are extracted and used to build *de-novo* assemblies using Trinity [30]. This module is useful to try to reconstruct and extend the sequence of an EVE that is not present in the reference assembly and can detect any lateral gene transfer event based on the list of non-host sequence used in the search.

~~~
$ bash ViR_LTFinder.sh \
-analysis_name SAMPLE_ID \
-read_list sample_reads.txt \
-working_dir working_directory/ \
-non_host_fasta non-host_sequence.fasta \
-trinity_exe Trinity
-th 32 -mem 64G \
~~~

#### 3.3.4 ViR_LTFinder.sh output

If there are reads supporting a novel integration, the script will produce a *de-novo* assembly created by Trinity using the consensus of the reads aligned to the sequence. Output of “ViR_LTFinder.sh” includes files for visualization of the aligned reads using a genomic data visualizer (e.g., IGV). Please carefully check the reads supporting the assemblies because multiple, or mosaic, assemblies may be built by Trinity, especially in case of viral integrations occurring in highly repetitive regions such as piRNA clusters.

## NOTES

1 Even though we suggest using a HPC cluster or a server, this protocol could run on a Unix based personal computer. The EVEs annotation (procedure 1) time-limiting commands are the BLASTx searches, which can be run on a personal computer with at least a quad-core CPU and 8GB of RAM, provided that the genome and the database are not too large. Procedures 2 and 3 could be run on a PC in case of a small genome assembly (<0.5Gb) and limited size fastq files, but even in that case we suggest running them on a workstation with at least 8 threads.

2 Both the SVD and Vy-PER [12] pipelines use BWA [14] to align WGS reads to a reference genome. Nevertheless, while the SVD pipeline uses BWA mem (which is now considered the standard BWA mapping program), Vy-PER was designed to run with sam alignments produced by BWA aln and sampe. We never tested Vy-PER with alignments produced with BWA mem but it is theoretically possible to use these data with Vy-PER. This possibility should be considered if WGS alignments done with BWA mem are already available or have been produced for the SVD pipeline.

## REFERENCES

1. Soucy SM, Huang J, Gogarten JP (2015) Horizontal gene transfer: building the web of life. Nat Rev Genet 16:472–482. https://doi.org/10.1038/nrg3962

2. Keeling PJ, Palmer JD (2008) Horizontal gene transfer in eukaryotic evolution. Nat Rev Genet 9:605–618. https://doi.org/10.1038/nrg2386

3. Chen Y, Williams V, Filippova M, et al (2014) Viral Carcinogenesis: Factors Inducing DNA Damage and Virus Integration. Cancers (Basel) 6:2155–2186. https://doi.org/10.3390/cancers6042155

4. Frank JA, Feschotte C (2017) Co-option of endogenous viral sequences for host cell function. Curr Opin Virol 25:81–89. https://doi.org/10.1016/j.coviro.2017.07.021

5. Dheilly NM, Adema C, Raftos DA, et al (2014) No more non-model species: the promise of next generation sequencing for comparative immunology. Dev Comp Immunol 45:56–66. https://doi.org/10.1016/j.dci.2014.01.022

6. Blair CD, Olson KE, Bonizzoni M (2020) The Widespread Occurrence and Potential Biological Roles of Endogenous Viral Elements in Insect Genomes. Curr Issues Mol Biol 34:13–30. https://doi.org/10.21775/cimb.034.013

7. ter Horst AM, Nigg JC, Dekker FM, Falk BW (2019) Endogenous Viral Elements Are Widespread in Arthropod Genomes and Commonly Give Rise to PIWI-Interacting RNAs. J Virol 93:e02124–18. https://doi.org/10.1128/JVI.02124-18

8. Kryukov K, Ueda MT, Imanishi T, Nakagawa S (2019) Systematic survey of non-retroviral virus-like elements in eukaryotic genomes. Virus Res 262:30–36. https://doi.org/10.1016/j.virusres.2018.02.002

9. Horie M, Honda T, Suzuki Y, et al (2010) Endogenous non-retroviral RNA virus elements in mammalian genomes. Nature 463:84–87. https://doi.org/10.1038/nature08695

10. Palatini U, Miesen P, Carballar-Lejarazu R, et al (2017) Comparative genomics shows that viral integrations are abundant and express piRNAs in the arboviral vectors *Aedes aegypti* and *Aedes albopictus*. BMC Genomics 18:1–15. https://doi.org/10.1186/s12864-017-3903-3

11. Altschul SF, Gish W, Miller W, et al (1990) Basic local alignment search tool. J Mol Biol 215:403–410. https://doi.org/10.1016/S0022-2836(05)80360-2

12. Forster M, Szymczak S, Ellinghaus D, et al (2015) Vy-PER: eliminating false positive detection of virus integration events in next generation sequencing data. Sci Rep 5:11534. https://doi.org/10.1038/srep11534

13. Pischedda E, Crava C, Carlassara M, et al (2021) ViR: a tool to solve intrasample variability in the prediction of viral integration sites using whole genome sequencing data. BMC Bioinformatics 1–15. https://doi.org/10.1186/s12859-021-03980-5

14. Li H, Durbin R (2009) Fast and accurate short read alignment with Burrows–Wheeler transform. Mass Genomics 25:1754–1760. https://doi.org/10.1093/bioinformatics/btp324

15. Robinson JT, Thorvaldsdóttir H, Winckler W, et al (2011) Integrative genomics viewer. Nat. Biotechnol. 29:24–26

16. Camacho C, Coulouris G, Avagyan V, et al (2009) BLAST+: architecture and applications. BMC Bioinformatics 10:421. https://doi.org/10.1186/1471-2105-10-421

17. Buchfink B, Xie C, Huson DH (2015) Fast and sensitive protein alignment using DIAMOND. Nat Methods 12:59–60. https://doi.org/10.1038/nmeth.3176

18. Whitfield ZJ, Dolan PT, Kunitomi M, et al (2017) The Diversity, Structure, and Function of Heritable Adaptive Immunity Sequences in the *Aedes aegypti* Genome. Curr Biol 27:3511–3519.e7. https://doi.org/10.1016/j.cub.2017.09.067

19. Quinlan AR, Hall IM (2010) BEDTools: a flexible suite of utilities for comparing genomic features. Bioinformatics 26:841–842. https://doi.org/10.1093/bioinformatics/btq033

20. Kitson E, Suttle CA (2019) VHost-Classifier: virus-host classification using natural language processing. Bioinformatics 35:3867–3869. https://doi.org/10.1093/bioinformatics/btz151

21. Shen W, Xiong J (2019) TaxonKit: a cross-platform and efficient NCBI taxonomy toolkit. bioRXiv

22. McKenna A, Hanna M, Banks E, et al (2010) The Genome Analysis Toolkit: a MapReduce framework for analyzing next-generation DNA sequencing data. Genome Res 20:1297–1303. https://doi.org/10.1101/gr.107524.110

23. Rimmer A, Phan H, Mathieson I, et al (2014) Integrating mapping-, assembly- and haplotype-based approaches for calling variants in clinical sequencing applications. Nat Genet 46:912–918. https://doi.org/10.1038/ng.3036

24. Garrison E, Marth G (2012) Haplotype-based variant detection from short-read sequencing. https://doi.org/arXiv:1207.3907[q-bio.GN]

25. Lai Z, Markovets A, Ahdesmaki M, et al (2016) VarDict: A novel and versatile variant caller for next-generation sequencing in cancer research. Nucleic Acids Res 44:1–11. https://doi.org/10.1093/nar/gkw227

26. Danecek P, McCarthy SA (2017) BCFtools/csq: haplotype-aware variant consequences. Bioinformatics 33:2037–2039. https://doi.org/10.1093/bioinformatics/btx100

27. Pischedda E, Scolari F, Valerio F, et al (2019) Insights Into an Unexplored Component of the Mosquito Repeatome: Distribution and Variability of Viral Sequences Integrated Into the Genome of the Arboviral Vector *Aedes albopictus*. Front Genet 10:93. https://doi.org/10.3389/fgene.2019.00093

28. Kent JK (2002) BLAT--The BLAST-Like Alignment Tool. Genome Res 12:656–664. https://doi.org/10.1101/gr.229202

29. Li H, Handsaker B, Wysoker A, et al (2009) The Sequence Alignment/Map format and SAMtools. Bioinformatics 25:2078–2079. https://doi.org/10.1093/bioinformatics/btp352

30. Grabherr MG, Haas BJ, Yassour M, et al (2011) Full-length transcriptome assembly from RNA-Seq data without a reference genome. Nat Biotechnol 29:644–652. https://doi.org/10.1038/nbt.1883

31. Chen S, Senar MA (2019) Exploring efficient data parallelism for genome read mapping on multicore and manycore architectures. Parallel Comput 87:11–24. https://doi.org/10.1016/j.parco.2019.04.014

32. Kondo H, Hirano S, Chiba S, et al (2013) Characterization of burdock mottle virus, a novel member of the genus Benyvirus, and the identification of benyvirus-related sequences in the plant and insect genomes. Virus Res 177:75–86. https://doi.org/10.1016/j.virusres.2013.07.015

33. Aguiar ERGR, de Almeida JPP, Queiroz LR, et al (2020) A single unidirectional piRNA cluster similar to the flamenco locus is the major source of EVE-derived transcription and small RNAs in Aedes aegypti mosquitoes. RNA 26:581–594. https://doi.org/10.1261/rna.073965.119

34. Fort P, Albertini A, Van-Hua A, et al (2012) Fossil rhabdoviral sequences integrated into arthropod genomes: Ontogeny, evolution, and potential functionality. Mol Biol Evol 29:381–390. https://doi.org/10.1093/molbev/msr226

35. Katzourakis A, Gifford RJ (2010) Endogenous viral elements in animal genomes. PLoS Genet 6:. https://doi.org/10.1371/journal.pgen.1001191

36. Chen X, Kost J, Li D (2019) Comprehensive comparative analysis of methods and software for identifying viral integrations. Brief Bioinform 20:2088–2097. https://doi.org/10.1093/bib/bby070

37. Mayer C (2010) Phobos 3.3.11

38. Gafni E, Luquette LJ, Lancaster AK, et al (2014) COSMOS: Python library for massively parallel workflows. Bioinformatics 30:2956–2958. https://doi.org/10.1093/bioinformatics/btu385

39. Kämpf C, Specht M, Scholz A, et al (2019) uap: reproducible and robust HTS data analysis. BMC Bioinformatics 20:664. https://doi.org/10.1186/s12859-019-3219-1

40. Morandi E, Cereda M, Incarnato D, et al (2019) HaTSPiL: A modular pipeline for high-throughput sequencing data analysis. PLoS One 14:e0222512. https://doi.org/10.1371/journal.pone.0222512

41. Li ITS, Shum W, Truong K (2007) 160-fold acceleration of the Smith-Waterman algorithm using a field programmable gate array (FPGA). BMC Bioinformatics 8:185. https://doi.org/10.1186/1471-2105-8-185

